# Short Tandem Repeat Profiling via Next Generation Sequencing for Cell Line Authentication

**DOI:** 10.1101/2023.02.25.530013

**Authors:** Yi-Hsien Chen, Jon P. Connelly, Colin Florian, Xiaoxia Cui, Shondra M. Pruett-Miller

**Affiliations:** Genome Engineering & Stem Cell Center (GESC@MGI), Department of Genetics, Washington University School of Medicine, St. Louis, 63110, USA; St. Jude Children’s Research Hospital, Department of Cell & Molecular Biology, Memphis 38105, USA; Center for Advanced Genome Engineering

**Author notes:** These authors contributed equally to this work (Yi-Hsien Chen, Jon P. Connelly).

**Keywords:** next generation sequencing, NGS, short tandem repeat, STR, targeted deep sequencing, cell line authentication, capillary electrophoresis, cell identity

## Abstract

Cell lines are indispensable models for modern biomedical research. In the era of CRISPR gene editing, they serve as versatile tools for preclinical studies, allowing patient specific mutations to be modeled or corrected and the resulting phenotypic outcomes studied. A large part of their usefulness derives from the ability of a cell line to proliferate over multiple passages (often indefinitely) allowing multiple experiments to be performed. However, over time, the cell line identity and purity can be compromised by human errors. Both cross contamination from other cell lines and even complete misidentification are possible. Routine cell line authentication is a necessary preventive measure and has become a requirement for many funding applications and publications. Short tandem repeat (STR) profiling is the most common method for cell line authentication and is usually carried out using standard polymerase chain reaction (PCR)-capillary electrophoresis (CE) analysis (STR-CE). Here we evaluated next generation sequencing (NGS)-based STR profiling of human and mouse cell lines at 18 and 15 loci, respectively, in a high-throughput format. Using the program STRight written in Python, we demonstrate that NGS-based analysis (STR-NGS) is superior to standard STR-CE in terms of the ability to report the sequence context of repeat motifs, sensitivity, and flexible multiplexing capability. STR-NGS is a valuable alternative for cell line authentication.

## Introduction

Since the introduction of short tandem repeats (STRs) as polymorphic DNA signatures (Puers et al., 1993), STR profiling has become the gold standard for identity confirmation in contemporary forensic science (Butler, 2007). STRs, also known as microsatellites or simple sequence repeats, are DNA segments containing core repeat units of 2-6 nucleotides that are scattered throughout the genome (Ellegren, 2004). In addition to the original 13 core STR loci (Bruce Budowle, 1997), 7 more loci were included in the Combined DNA Index System (CODIS) that is used for forensics in the United States (Hares, 2015). These STR loci are highly polymorphic, genetically unlinked, and offer powerful and accurate individual identification.

Human and mouse cell lines are important research models for mechanistic studies, target identification and therapeutic development. However, cell cultures are at risk for misidentification due to human errors and cross-contamination from other cell lines. Examples of misidentified cell lines jeopardizing scientific research continue to grow in number (Nardone, 2007), demonstrating the urgent need for frequent cell line authentication. As a result, cell line authentication is now required by an increasing number of journals prior to publication as well as for grant applications (Almeida et al., 2016). A method that is sensitive, high-throughput and economical is highly desirable.

The framework of using STRs for the authentication of human cell lines was first introduced in 2010 (Barallon et al., 2010). To date, many thousands of human cell line STR profiles are available, reflecting the unique donors from whom they were originally derived (Novroski et al., 2016). The American Tissue Culture Collection (ATCC) has published standard guidelines which recommend the use of at least eight STR loci (TH01, TPOX, vWA, CSF1PO, D16S539, D7S820, D13S317 and D5S8181 plus Amelogenin for gender identification) for human cell line authentication (ASN-0002, 2011). Moreover, recent studies report several additional STR loci that may be used to authenticate mouse cell lines (Almeida et al., 2019; Almeida et al., 2014).

Currently, STR profiling is predominately performed by resolving multiplexed, fluorescently-labeled PCRs using capillary electrophoresis (Deforce et al., 1998). However, loci of the same size but with different sequences cannot be distinguished using this conventional method. The full sequences and nucleotide variations found in STR loci provide additional data that aid in identification. The conventional STR method also requires access to a specialized instrument, a Genetic Analyzer, which has limited potential for further improving sensitivity or throughput. With continuous technical improvements, NGS has become an attractive alternative for STR profiling. Different NGS platforms such as Roche/454, Ion torrent, and Illumina systems, have proven capable of sequencing the majority of STR loci in forensic science (Fordyce et al., 2015; Mikkelsen et al., 2014; Van Neste et al., 2014). NGS-based STR (STR-NGS) profiling has several advantages over the conventional method, including high-throughput, low cost when running many samples, flexibility with which STR loci to include, quantitative measurements for mixed samples, and high resolution for single nucleotide polymorphisms (SNPs) (Bornman et al., 2012; Shin et al., 2017). This method has yet to be applied to cell line authentication.

In the present study, we evaluate the accuracy and sensitivity of the Illumina MiSeq platform for STR profiling of human and mouse cell lines and demonstrate the method is valid and scalable for routine quality control of human and mouse lines used in biomedical research.

## Materials and methods

### Control samples and Cell lines

Two human iPSC lines, BJFF.6 from a male donor, and AN1.1 from a female donor were generated at the Genome Engineering & Stem Cell Center (GESC) at Washington University in St. Louis. iPSCs were maintained on Matrigel (Corning) coated plates in Stem-Flex medium (Thermo Fisher). In addition, four mouse cell lines, NIH3T3, MC3T3-E1, CT26 and 4T1 were obtained from the American Type Culture Collection (ATCC) and cultured in the ATCC-recommended media. All cell lines were grown in a humidified incubator set at 5% CO_2_ and 37°C.

### Extraction and quantification of DNA

DNA was extracted using a crude DNA extraction buffer (10mM Tris pH8.0, 2mM EDTA, 0.2% Triton X-100 and 200 µg/ml proteinase K) or DNA Blood Mini kit (Qiagen) following the manufacturer’s instructions. DNA was quantified using NanoDrop One spectrophotometer (Thermo Fisher).

### STR loci and primer design

Eighteen STR loci (CSF1PO, D13S317, D16S539, D18S51, D19S433, D2S1338, D21S11, D3S1358, D5S818, D7S820, D8S1179, FGA, PentaD, PentaE, TH01, TPOX, vWA and Amelogenin) recommended from the ATCC and the Combined DNA Index System (CODIS) were used for human cell line profiling. Fifteen STR loci (18-3, 4-2, 5-5, 6-7, 9-2, 12-1, 15-3, X-1, 1-1, 2-1, 3-2, 8-1, 11-2, 17-2 and 19-2) were used for mouse cell line STR profiling (Almeida et al., 2019; Almeida et al., 2014). The Primer Blast design tool from NCBI was used to design PCR primers flanking STR regions based on the reference sequences from STRBase (Ruitberg et al., 2001) (https://strbase.nist.gov/) for human loci and two recent studies (Almeida et al., 2019; Almeida et al., 2014) for mouse loci, respectively. To increase PCR specificity, nested forward primers were used for the PentaE locus and only the internal forward primer has the adapter sequence required for NGS. All primer sequences are listed in Table S1 and S2.

### Optimized PCR amplification and processing using Illumina chemistry

A two-step PCR strategy is used to amplify each STR locus for NGS. PCR1 amplifies the individual STR locus and adds a partial Illumina adapter. A portion of the PCR1 amplicon is then used as template for PCR2 and adds a unique index and the remaining Illumina adaptor to each sample. PCR conditions for STR loci (PCR1) have been optimized. Results for detailed optimizations are found in Supplemental Figure 1. Tetramethylammonium (TMA) oxalate solution was made by mixing a 2 to 1 molar ratio of aqueous TMA hydroxide (Sigma Aldrich) and ammonium oxalate monohydrate (Thermo Fisher), respectively (Markowitz, 1957). PCR reactions contain 0.5 mM of each primer, 70 ng of genomic DNA, 2 mM TMA oxalate, and 1X Platinum SuperFi PCR Master Mix (Thermo Fisher). The samples were amplified using a Veriti thermal cycler (Thermo Fisher) as follows: 98°C for 2 min; 28 cycles of 98°C for 15 s, 60°C for 15 s, and then 68°C for 30 s; followed by an extension for 2 min at 72°C. For multiplex PCRs, samples were amplified using the following reaction mixture: 0.2 µM final concentration of primer sets for each locus (6 sets in each reaction), 140 ng of genomic DNA in the reaction, and 1X Platinum SuperFi PCR Master Mix (Thermo Fisher). Multiplex PCRs were carried out in a Veriti thermal cycler (Thermo Fisher) with the following cycle conditions: 98°C for 2 min; 30 cycles of 98°C for 15 s, 60°C for 45 s, and then 68°C for 1 min; followed by an extension for 5 min at 72°C. Products of PCR1 were subjected to an additional 5 cycles with non-overlapping dual-indexing primers without TMA oxalate, and products of PCR2 were pooled and purified using 0.6X SPRI beads. The purified library was run on a BioAnalyzer using a high sensitivity chip to confirm quality and quantity. It was then denatured with 0.1nM NaOH, followed by neutralization with 0.1M HCl. The denatured pool was diluted to 20pM following Illumina guidelines. It was sequenced on a MiSeqv2 using a MiSeq Reagent Kit 500v2 kit with the following inputs: 250 bp Read1, 10 bp Index1, 8 bp Index2 and 250 bp Read2. Samples were demultiplexed using the index sequences, FASTQ files were generated using the Illumina bcl2fastq software, allowing 1 bp mismatch.

**Figure 1.**
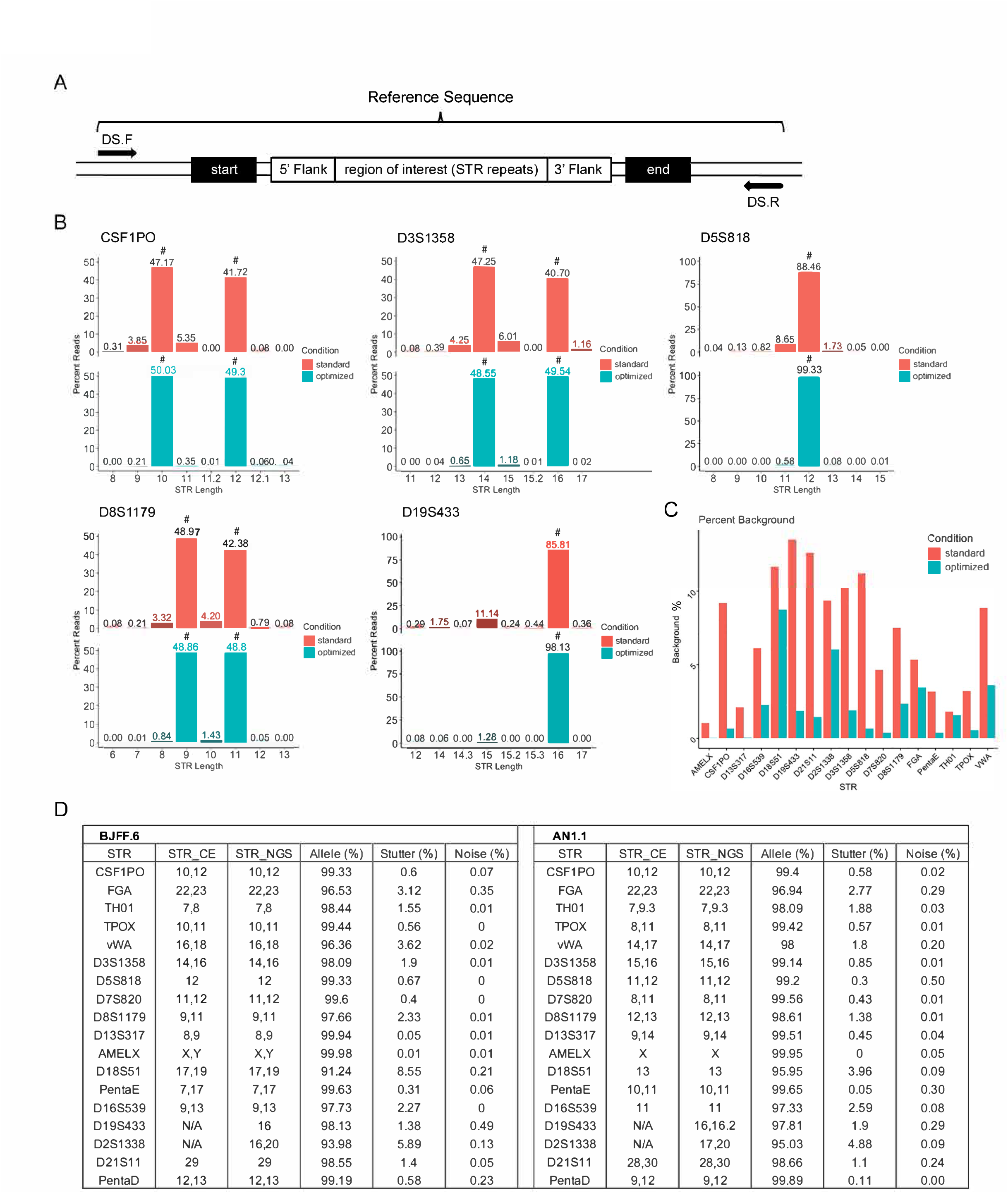
STR-NGS optimization and performance. (A) Schematic representation of a STR locus with STR repeats, flanking regions at the start and end of the repeat region, and a targeted deep sequencing (DS) primer pair. Sequence from “5’ Flank” to “3’ Flank” is used for repeat length analysis. (B) Allele and stutter frequencies for five representative STR loci calculated as the percentage of the parent allele reads. Verified STRs indicated by a # above bar plot. (C) Total background (stutter + noise) in all STR loci in the comparison between standard and optimized PCR conditions. (D) STR profile for each locus examined by CE and NGS based methods in two diploid iPSC lines.

### STR data analysis

Human STR-CE profiling was performed by Cell Line Genetics (WI, USA). STR-NGS was analyzed using the python script STRight. This program is a modified version of CRIS.py (Connelly and Pruett-Miller, 2019) and takes a csv file containing target STR data as input. The csv files (STR_human.csv and STR_mouse.csv) contain 9 columns: (1) STR name, (2) Reference sequence, (3) Start target sequence, (4) End target sequence, (5) repeat size, (6) bp_modifier, (7) the number of repeats in a reference sequence found on strbase.nist.gov (this can be used as a quality control check if novel STRs are being added), (8) SNP_modifier, and (9) notes. Briefly, the program runs through each line of the fastq files and tests whether the start and end target sequences of each STR can be matched to the read. If both sequences are found, the distance between the two sequences is measured in base pairs. Because SNPs could occur in the flank region outside the STR repeat region, the start and end sequences for STRight analysis do not always land precisely at the beginning and end of each locus. Instead, they may be placed several base pairs upstream or downstream of the repeat to avoid known SNPs. In addition, some STR loci have additional nucleotides inside the repeats (i.e., DS21S11 has 13 additional nucleotides among “TCTA” and “TCTG” repeats). To take these extra base pairs into account, the bp_modifier value is used to subtract any extra base pairs from the total length (i.e., if the start sequence lands 5bp upstream from the start of the STR repeat region, then a bp_modifier of 5 would subtract 5 base pairs from the measured distance). The repeat number of a locus is determined by dividing the calculated distance by the size of each repeat, for example, 4 for an “AAAG” repeat. Finally, there are special cases where SNPs in a given locus create an extra repeat. For example, the STR D13S317 is a “TATC” repeat whose end usually terminates right before the sequence “AATC”. Some individuals however have an A>T variant creating a termination sequence of “TATC”, and which also adds an additional repeat. Typical STR-CE utilizes a primer mixture containing two reverse primers which have the 5’ nucleotide landing on either the “A” or “T” of the SNP to detect that variant. STRight contains a SNP_modifier which checks whether the A/T SNP is present in the read. If the indicated SNP (in this case “T”) is present, then the SNP_modifier value (in this case 1), would add an extra repeat to the total number to account for the SNP. STRight reports the top repeat lengths for each STR along with sequence information. STRight is free to use and source code is available at https://github.com/patrickc01/STRight

### Analysis of mixed samples

Genomic DNA was extracted from diploid iPSC cell lines, BJFF.6 and AN1.1 cells, and added to individual reactions with a final concentration of 140 ng of total DNA at BJFF.6/AN1.1 ratios of 1:1, 5:1, 10:1, 20:1, 100:1, 200:1, and 1000:1. PCR amplification is described above, and analysis was performed in the same way as the single source samples.

## Results

### NGS optimization for human STR profiles

We developed an NGS-based STR profiling (STR-NGS) method that amplifies and analyzes common human and mouse STR loci for repeat length. Primers designed to each STR locus amplify a region of interest (ROI) containing the STR repeats flanked by left and right consensus sequences (Figure 1A). Our NGS library construction is a two-step process. PCR1 amplifies the region of interest and adds partial Illumina adaptors to the amplicons. PCR2 is the indexing PCR and adds unique identifiers to each sample. To improve NGS read quality, we optimized the PCR1 reaction at each locus. First, different primer sets were tested for amplification efficiency and specificity, and the final primer pairs for the 17 CODIS STR loci and sex (AMEL) locus in this study are listed in Table S1. Second, we evaluated different DNA polymerases, including several high-fidelity DNA polymerases-AccuPrime Taq DNA polymerase high fidelity from Thermo Fisher, Platinum SuperFi PCR Master Mix from Thermo Fisher, and Titanium Taq DNA polymerase from Takara. MyTaq from Bioline is a low cost and high-performance polymerase and was used as a standard for comparison. We also tested and optimized the quantity of genomic DNA sample input. We found that Platinum SuperFi PCR mix produced higher PCR amplicon yield and superior accuracy for this application (data not shown). SuperFi was used for subsequent tests and studies. As reported previously, we found that the quantity of input template DNA has a substantial effect on the outcome of PCR reactions (Lorenz, 2012). The optimal DNA amount in our assays is between 70-140 ng per reaction, roughly equivalent to 10-20,000 cells (Figure S1A and B). Low amplification was observed with input cell numbers lower than this range (data not shown). PCR supplements have been shown to be effective in improving yields of specific PCR products at difficult loci (Lorenz, 2012). Given the presence of homopolymer stretches at many STR loci (Gettings et al., 2015), poor PCR amplification can result in higher error rates in NGS. Several tetramethylammonium (TMA) derivatives have been shown to increase the specificity of PCR and improve the yield of amplification products (DiLella and Woo, 1987; Kovarova and Draber, 2000). We tested two TMA derivatives, TMA chloride (TMAC) and TMA oxalate (TMAO). TMAC addition in PCRs made a modest improvement on reducing background in most STR loci compared to PCR with no additive (data not shown), while addition of TMAO to PCR reactions increased the yield of specific PCR products (Figure S1A and B). Therefore, SuperFi plus TMAO was used for PCR1 in subsequent studies.

We also optimized the indexing reaction (PCR2). Dual-indexing has proven to increase the accuracy of multiplex sequencing and throughput on the NGS platform (Kircher et al., 2012). High-fidelity polymerases, such as SuperFi, are effective at reducing errors introduced in the indexing reaction (Figure S1B). Five representative STR loci are highlighted in comparison between standard and improved PCR conditions (Figure 1B) as well as the overall background improvement in all loci (Figure 1C). Moreover, it should be noted that amplification efficiency varies at different loci, as measured by total NGS reads at each locus (Figure S1C).

### STR-NGS vs. STR-CE

We next compared the results of STR-NGS using our optimized conditions to STR-CE conducted by Cell Line Genetics (Madison, WI) for two diploid iPSC lines. The two sets of data are highly in agreement (Figure 1D). In both cell lines, noise, stutter, and allele percentages are shown for all 18 STR loci (Figure 1D). Stutters are defined as sequence reads that contain a number of STR repeat lengths that are smaller or larger (predominantly 1 repeat shorter) than the actual allele and with a frequency of less than 10%. Noise is defined as reads that are not alleles or stutters and are likely the product of sequencing errors or off-target amplification. Among all STR loci, 91-99% of the reads from STR-NGS corresponded to parent alleles and noise comprised less than 1% of the total reads. The stutters ranged from 0 to 3.6%, except D18S51 and D2S1338 which had higher stutter percentages. Together, we demonstrate that STR-NGS identifies human cell lines as accurately as the standard STR-CE method.

### Sensitivity and multiplexing capability of STR-NGS

Early detection of cross contamination in a given culture is an important application of STR-NGS. We evaluated STR-NGS sensitivity in detecting a contaminating cell line sample among varying mixed ratios of BJFF.6 and AN1.1 cells, where AN1.1 represented the minor component of the mixtures. These two lines share STRs of the same length at several loci, therefore we focused on unique repeats found in AN1.1. We found that the AN1.1 fractions by read count correlate well with the expected ratios (Figure 2A). Next, we examined the D81179 locus specifically for its distinct repeat lengths in two cell lines. Given the major stutter of D81179 is 4 bp shorter than the parent allele (Figure 1B), it provided a good opportunity to identify the minor alleles from AN1.1. We found STR-NGS could detect a minor contamination level as low as 0.5% (1:200) within a sample (Figure 2B). Note the AN1.1 contaminating mixture has STR repeat lengths that are 1-2 repeats longer than the BJFF.6 repeats which allow them to be differentiated from stutters.

**Figure 2.**
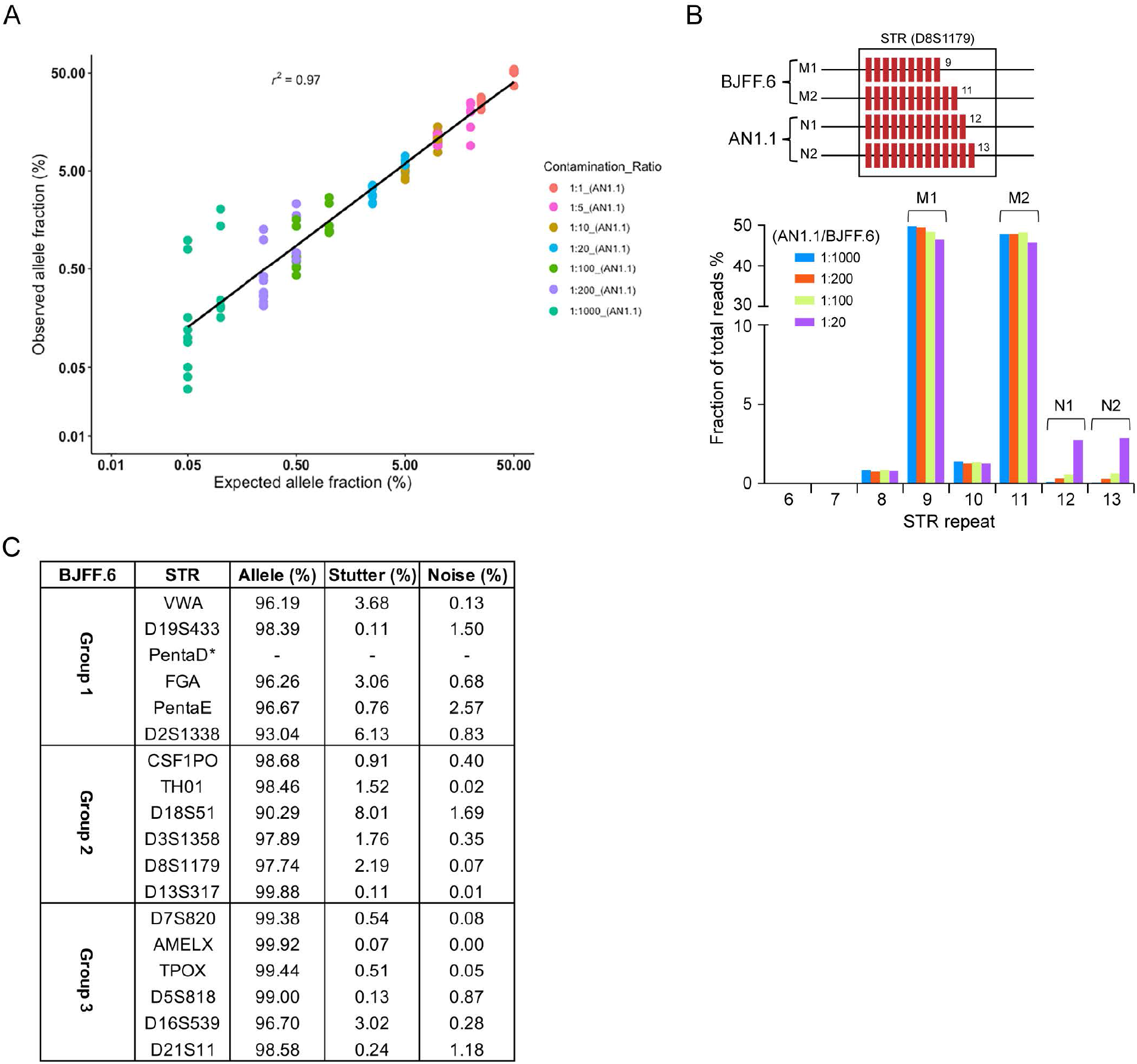
STR-NGS sensitivity on a mixed sample and optimized multiplexed conditions. (A) Observed allele fractions of informative STRs repeats are plotted against the expected ratio for given mixtures of 2 genomic samples. AN1.1 gDNA was diluted into BJFF.6 gDNA in a ratio of 1:1 to 1:1000. Expected allele fractions for diluted AN1.1 STRs correlate well with observed. (B) A mixture of two diploid cell lines was analyzed for STR D8S1179. M and N alleles indicate genotypes from the major and minor components, respectively. Bar graph shows percentage of read counts for each STR repeat of both cell lines in different mixture ratios. (C) STR profile for each locus examined using multiplexed STR-NGS in BJFF.6 iPSC cells. (*) dropout allele.

To reduce the cost and labor of the STR-NGS assay, we explored the feasibility of multiplexing the PCR reactions. We grouped different numbers of STR loci for multiplexed PCRs. Up to six loci can be amplified in one PCR reaction without compromising stutter and noise levels (Figure 2C). Stutter levels for D18S51 and D2S1338 remained higher than other loci regardless of PCR conditions, and dropout of the PentaD STR was observed during multiplexing reactions. Overall, we were able to obtain satisfactory percentages of clean reads for all loci except for PentaD and conclude that STR-NGS is reliable in multiplexed format.

### STR-NGS in mouse cell line authentication

With great effort from the scientific community, STRs have been identified that allow authentication of mouse cell lines from different mouse strains (Almeida et al., 2019; Almeida et al., 2014). To determine whether STR-NGS could be used for mouse cell line authentication, we performed STR-NGS on two previously reported mouse cell lines, MC3T3-E1 and NIH3T3 derived from different mouse strains (Almeida et al., 2014). First, both cell lines were used to optimize PCR performance and optimized primer sets are listed in Table S2. For MC3T3-E1 all loci matched the published STR-CE results. Of note, the STRs at loci 6-7, and 15-3, had higher stutter or noise percentages than the other STRs assayed (Figure 3A). Nonetheless, the combined stutter and noise for all STRs for MC3T3-E1 was less than 5% of the total reads. However, for NIH3T3 cells, minor fractions of different repeat lengths were observed at loci 4-2 and 18-3 (Figure 3A and 3B).

**Figure 3.**
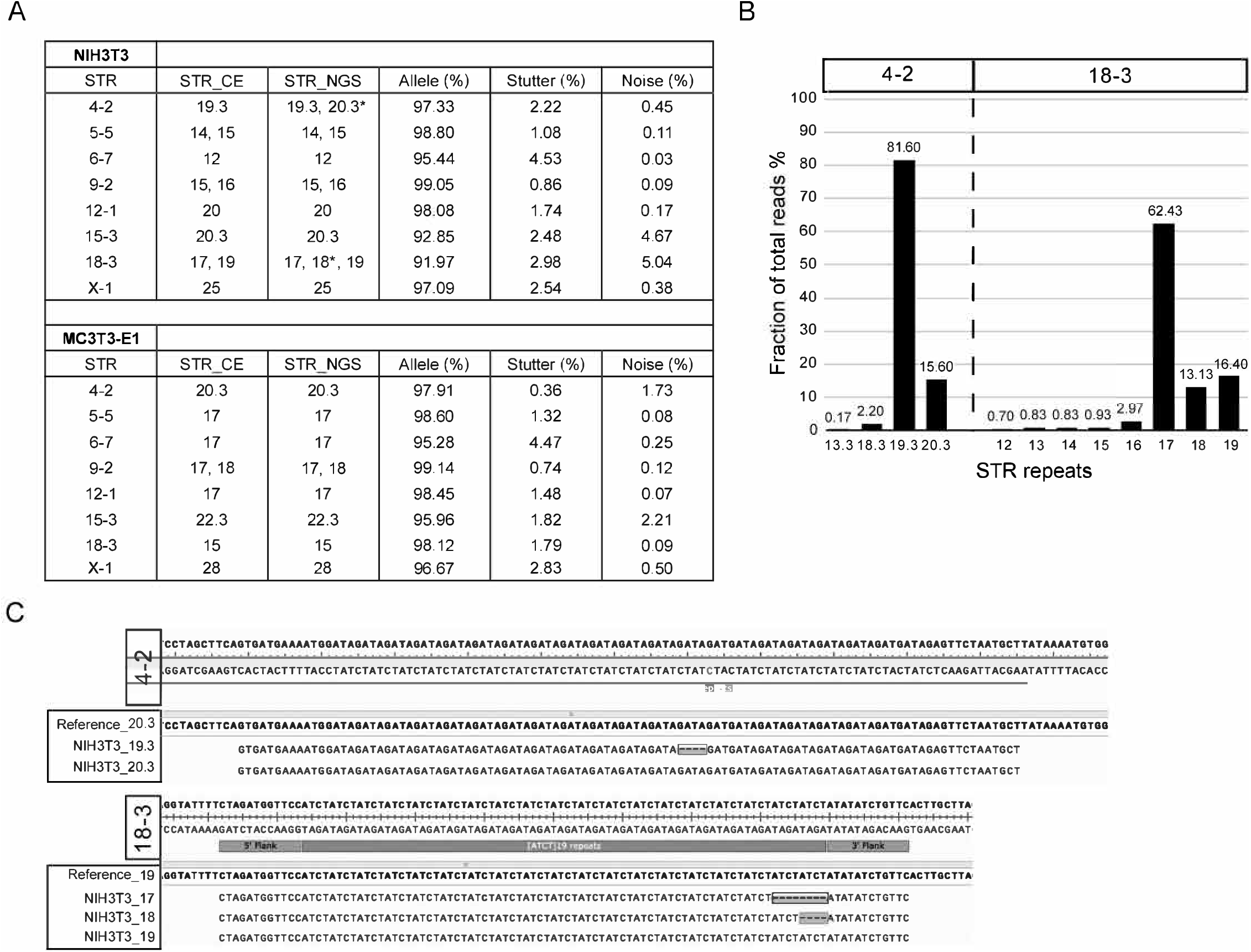
STR-NGS in mouse cell lines. (A) STR profile in two mouse cell lines (NIH3T3 and MC3T3-E1) shows allele, stutter and noise fractions of total parent allele reads. Asterisk indicates the difference in STR repeat length between reference and STR-NGS. (B) The frequencies of different stutters and repeat lengths, and (C) sequence context of different STR repeat lengths in two STR loci (4-2 and 18-3) different between reference and STR-NGS in NIH3T3 cells.

To investigate the reason for the discrepancy in STR repeat length in the NIH3T3 cells, we compared the read percentages and sequencing content of different repeat lengths in STR loci, 4-2 and 18-3. Stutter products in most loci usually comprise less than 5% of total reads in both human and mouse samples (Figure 1C and 3A). The read percentages for additional lengths were 15.60% for 4-2 and 13.13% and 16.40% for 18-3, which is unlikely the result of suboptimal assay conditions. One of the main advantages of the NGS based method is that the raw data contains the actual sequence of a given locus. We examined the STR-containing reads with complete microsatellite sequences. At STR 4-2, our finding is consistent with results from a recent study (Almeida et al., 2019) which show 19.3 and 20.3 STR repeats for NIH3T3 cells (Figure 3C). At STR 18-3, we observed reads corresponding to all three repeat lengths – 17, 18, and 19. We hypothesized that the heterogeneity of the immortalized NIH3T3 cells might have another population harboring 17 and 18 repeats. To verify the 18 repeats did not come from stutter products of 17 and 19, we performed single cell sorting and analyzed single cell clones from parental NIH3T3 cells. Three out of the five independent clones carried 17 and 18 repeats at the locus. (Table 1 and Figure S2). Another advantage of STR-NGS is that some information can be inferred based on the number of sequence reads for each repeat length. For example, all five clones show an allelic ratio of approximately 2:1 for either the 17:18 or 17:19 repeats (Figure S2), which suggests that there are three copies of this locus.

**Table 1.**
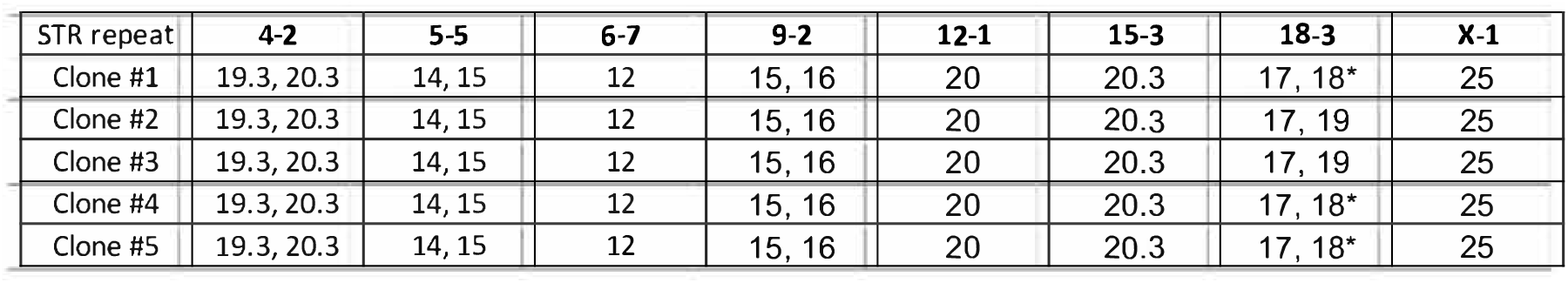
STR profiles of five NIH3T3 clones using STR-NGS (*) difference in repeat length between STR-NGS and STR-CE method

| STR repeat | 4-2 | 5-5 | 6-7 | 9-2 | 12-1 | 15-3 | 18-3 | X-1 |
| --- | --- | --- | --- | --- | --- | --- | --- | --- |
| Clone #1 | 19.3, 20.3 | 14, 15 | 12 | 15, 16 | 20 | 20.3 | 17, 18* | 25 |
| Clone #2 | 19.3, 20.3 | 14, 15 | 12 | 15, 16 | 20 | 20.3 | 17, 19 | 25 |
| Clone #3 | 19.3, 20.3 | 14, 15 | 12 | 15, 16 | 20 | 20.3 | 17, 19 | 25 |
| Clone #4 | 19.3, 20.3 | 14, 15 | 12 | 15, 16 | 20 | 20.3 | 17, 18* | 25 |
| Clone #5 | 19.3, 20.3 | 14, 15 | 12 | 15, 16 | 20 | 20.3 | 17, 18* | 25 |

Next, we expanded the number of total mouse STR loci to 15 based on a recent report (Almeida et al., 2019) and compared the results between STR-CE and STR-NGS using three different mouse cell lines – NIH3T3, CT26, and 4T1 used in that study. In all three lines, STR-CE and STR-NGS results matched at 12 of 15 loci tested (Table 2). To assess the differences among the three cell lines, we performed sequencing alignment with the data from parental cells and single cell clones. In NIH3T3 cells, STR-NGS was able to detect the alleles from different subpopulations present in the pool (Figure S3). Three unique subpopulations were identified by the presence of three different repeat lengths in STR locus 11-2 (Figure S3A). To investigate further, we isolated, expanded, and analyzed 12 single cell clones. Most clones were represented by repeat lengths of 15 and 17. However, five of twelve clones presented with 15,16,17 or 14,15,17 repeat lengths (Figure S3B). A unique clonal population was also observed in 4T1 cells (Figure S4) where the 17-2 locus differed between STR-NGS and STR-CE. STR-CE only shows a repeat length of 15 in 4T1 cells compared to repeats of 14 and 15 given by STR-NGS (Table 2). Given that the stutter for 17-2 locus was below 6% in NIH3T3 and CT26 cell lines (Table 2), we believe a repeat of length 14 exists and is from the minor allele contributor (Figure S4B). Together, these data demonstrate that STR-NGS offers great accuracy on allele calling compared to the traditional STR-CE method and is a valuable alternative for mouse cell line authentication.

**Table 2.**
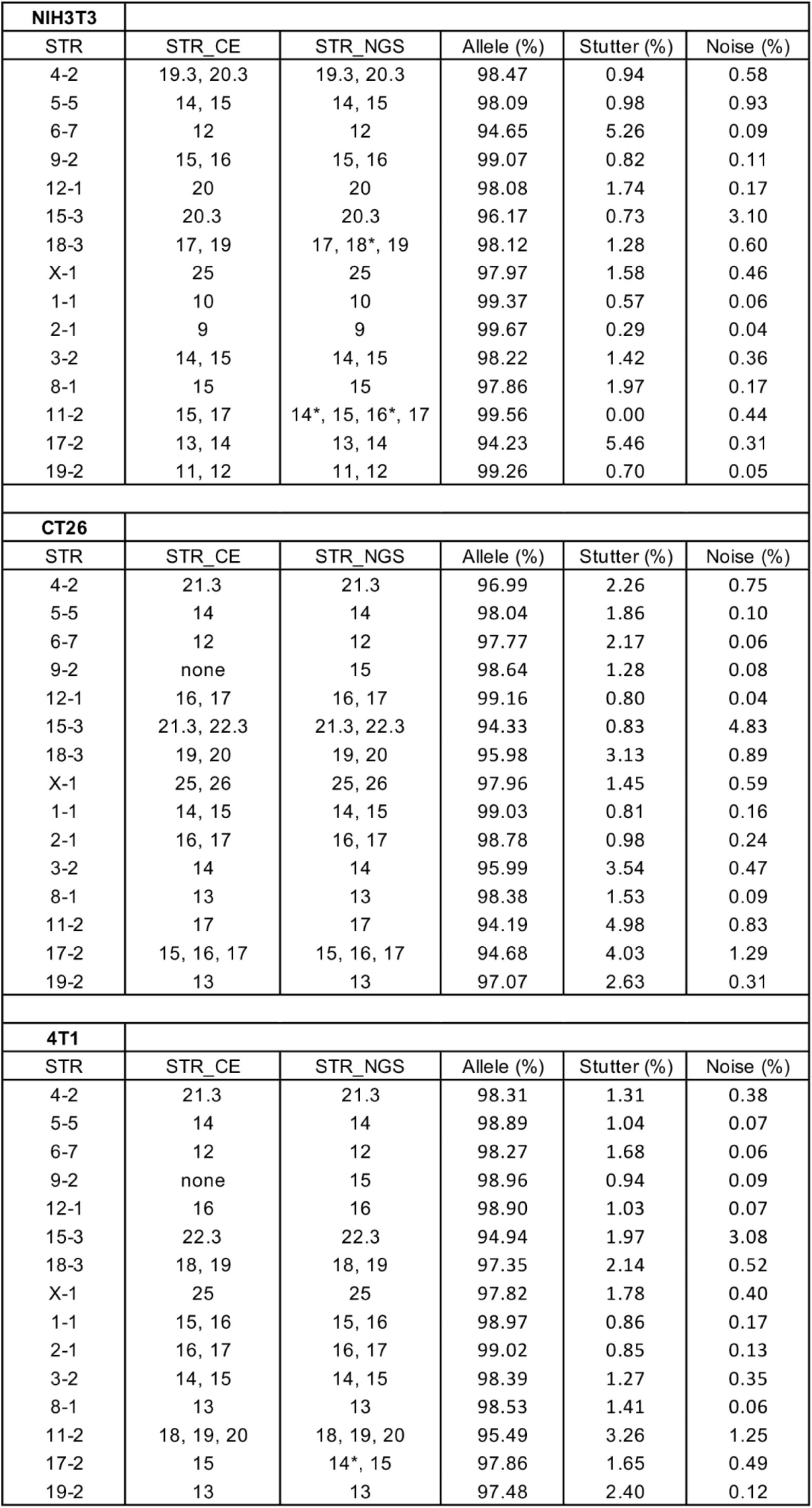
STR profiles of 3 mouse cell lines (*) difference in repeat length between STR-NGS and STR-CE method

## Discussion

STR profiling is the gold standard for human identity testing (Butler, 2007) and cell line authentication (Barallon et al., 2010), given the uniqueness of each individual and relatively easy assay format. STR-CE is widely used for separating STR markers based on size distribution; however, it has limited sensitivity and does not provide the STR sequence identity or context. Although NGS has been implemented for human identification in forensic genetics (Børsting and Morling, 2015)., cell line authentication has been limited to STR-CE. Because of heterogeneity within many cell lines, this lack of sensitivity may lead to cell line misidentification or delayed identification of cross contamination. Here, we present a simple and robust NGS-based solution for human and mouse cell line authentication using optimized amplification conditions for each STR loci and a novel STR analysis program called STRight.

To develop a robust STR-NGS workflow, we first evaluated different DNA polymerases, cell number, indexing PCR conditions, and additives on STR loci amplification. After optimization, we achieved over 96% allele specific signals at most loci. Next, we compared the results from STR-CE and STR-NGS workflows on two diploid human iPSC lines. Overall, the STR-NGS and STR-CE results matched. Moreover, except for D18S51 and D2S1338, the average range of stutter and noise percentage observed per STR locus is below 5% and 1%, respectively, for STR-NGS compared to the stutter ratio of 20% for STR-CE which is often used for initial filtering (Brookes et al., 2012; Mikkelsen et al., 2014; Novroski et al., 2016). While sequencing errors were very minimal in most STR loci, D18S51 and D2S1338 are obvious underperformers in the current protocol with a higher proportion of reads containing errors in the ROI (data not shown). It is unclear at which point the errors are introduced, but a longer ROI and sequence context could be potential factors contributing to the stutter percentage and sequencing errors.

A previous study showed that in mixed samples, sequences from the minor fraction (down to 1%) were detectable by NGS (Fordyce et al., 2015). In contrast, STR-CE has a general detection limit for the minor contributor in 1:10 mixtures (Westen et al., 2009). In this study, we show that STR-NGS is sensitive in detecting a minor contributor among a series of DNA mixtures down to 0.5% (Figure 2B).

Multiplexed PCRs for STR typing are both time and cost-effective. Indeed, STR alleles are routinely analyzed by multiplexed PCRs followed by CE-based analysis (Butler, 2006, 2007). However, large variations in target loci lengths result in a bias towards loci with shorter amplicons and lead to an imbalanced locus coverage (Van Neste et al., 2012). While up to four or ten multiplexed PCRs for STR sequencing have been demonstrated for 454 or Ion Torrent sequencing platforms, respectively, high noise ratios and poor performance on allele calling have limited its utility (Butler, 2007; Fordyce et al., 2015). In this report, we demonstrate that our optimized workflow for STR-NGS with TMAO additive is capable of multiplexing 6 STR loci with less than 5% stutters of total reads in 15 of 17 total human STR loci. Moreover, we recommend that underperforming loci, such as PentaD, be avoided in a multiplexed PCR setting. A careful selection of loci and further PCR multiplex optimization could further improve the utility of the STR-NGS approach.

Unlike STR forensic analysis, STR analysis is more complicated in human cell lines due to the heterogeneity and aneuploidy present in many immortalized and transformed lines. Indeed, STR profiles for a given human cell line may even vary from different sources (https://web.expasy.org/cellosaurus/), and genetic variation within cell lines has been well documented (Pourquier and Azzi, 2019). The need for cell line authentication in mouse cell lines has also risen in recent years (Almeida et al., 2019; Almeida et al., 2014), so much so that the American Type Culture Collection is now offering mouse STR profiling as a service. In this study, we demonstrate for the first time to our knowledge that STR-NGS can be used in mouse cell line authentication. In addition to generating correct genotypes, STR-NGS data resulted in the identification of sub-repeat variants from sub-populations of NIH3T3 cells that were undetected by the STR-CE method (Almeida et al., 2019; Almeida et al., 2014). In contrast to fragment length analysis by STR-CE, sequencing compositions in the ROI by STR-NGS provides additional detail for better interpretation of STR variation.

## Conclusions

We developed a simple workflow that offers a reliable solution for genotyping the most frequently used STR loci in both human and mouse cell lines (Table 3 and 4) without the need for a specialized instrument made for STR. Multiplexing reactions provide the flexibility to run many samples at once and reduce both time and cost for the operation. STR-NGS improves allele calling efficiency, detection sensitivity, and allows better differentiation of artifacts from minor contributors than the standard STR-CE method. STR-NGS can be easily used to authenticate cell lines as well as sensitively detect cross contamination.

**Table 3.**
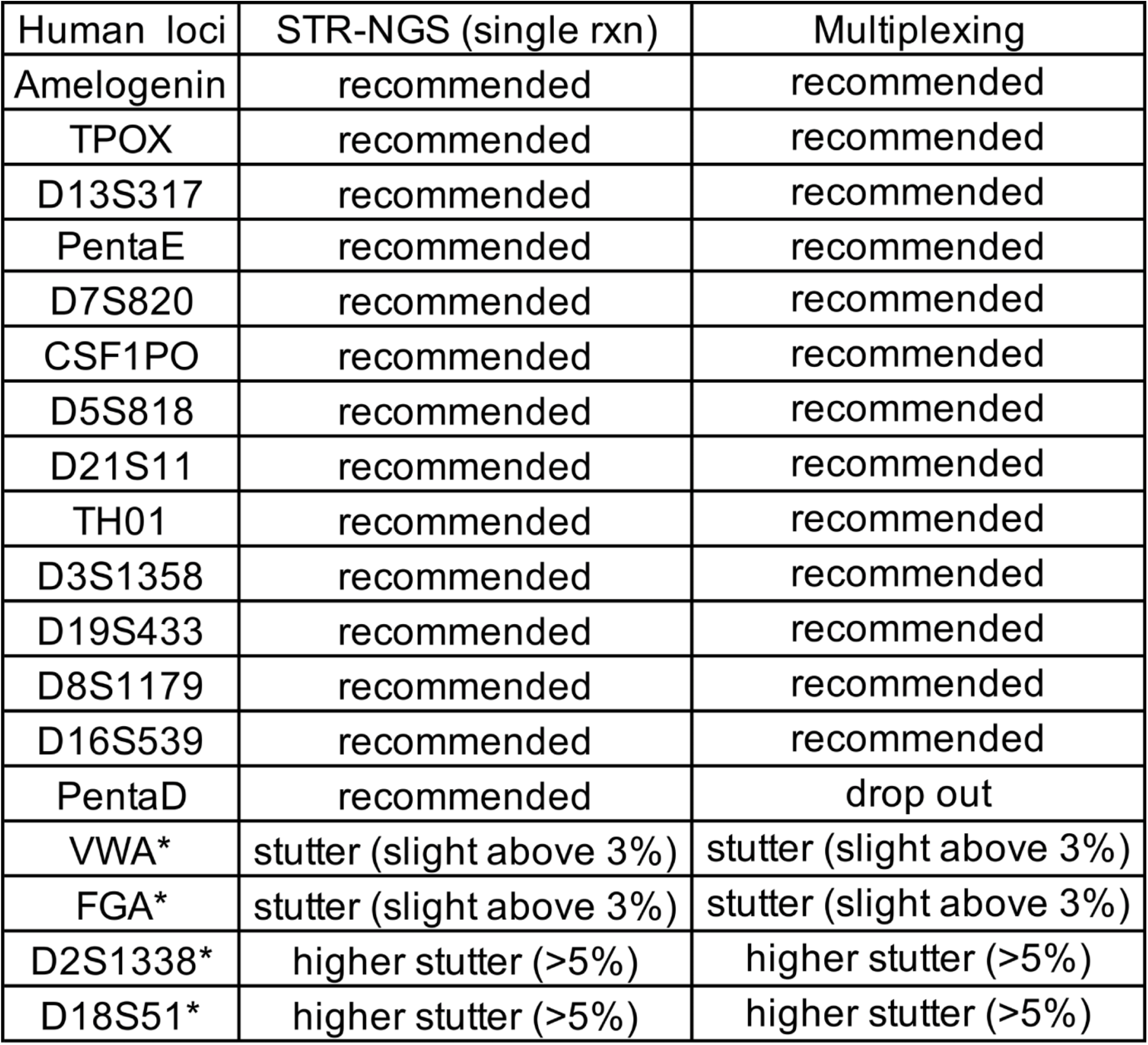
Summary of human STR loci using STR-NGS (*) Underperformer for human STR-NGS

**Table 4.**
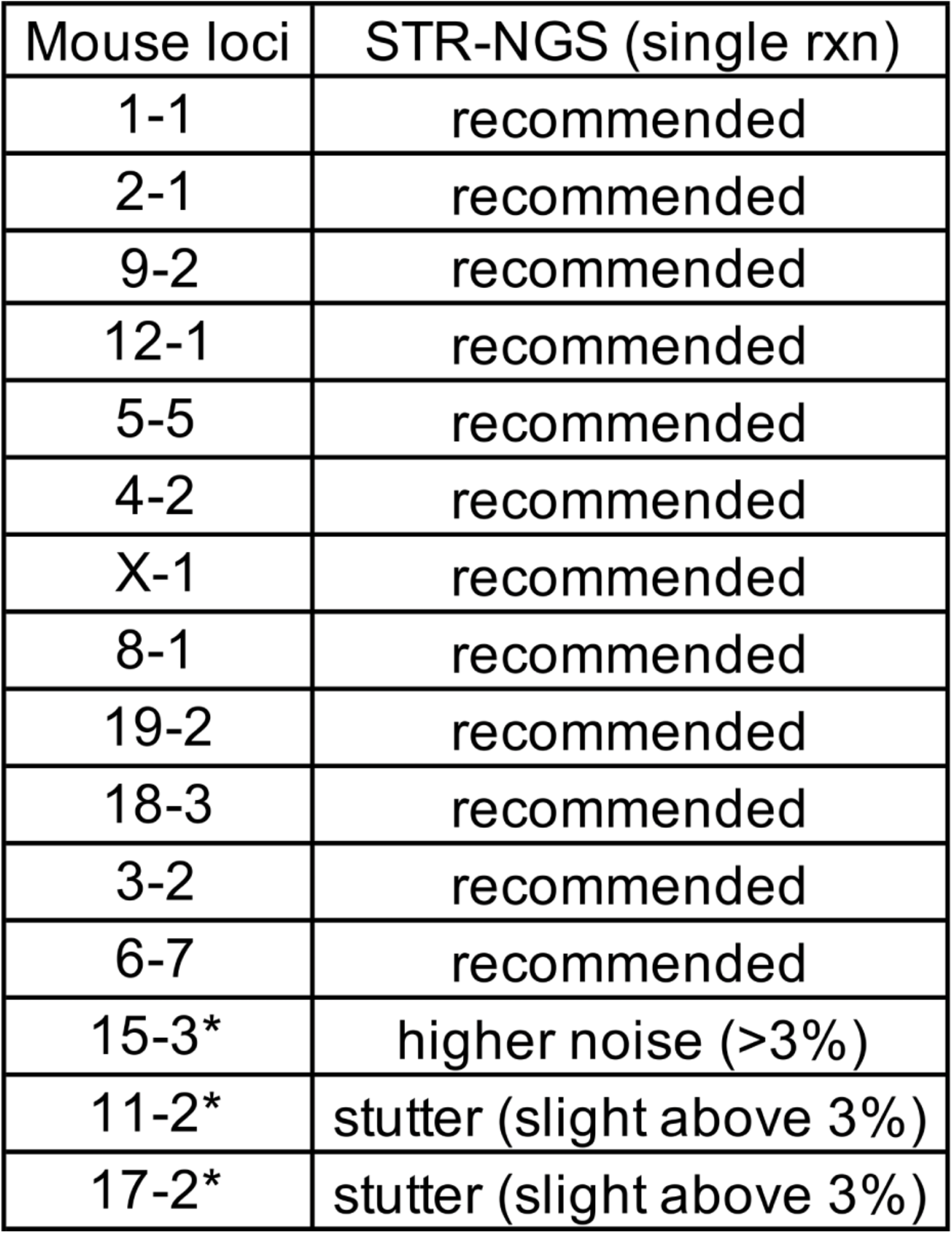
Summary of mouse STR loci using STR-NGS (*) Underperformer for mouse STR-NGS

## Acknowledgements

We thank Jane Kouranova for the initial primer design for mouse STR loci and Jed Lin for assisting NGS library preparation. We thank the DNA Sequencing Innovation Lab, The Edison Family Center for Genome Sciences & Systems Biology which provided NGS services.

## Figure Legends

**Figure S1.**
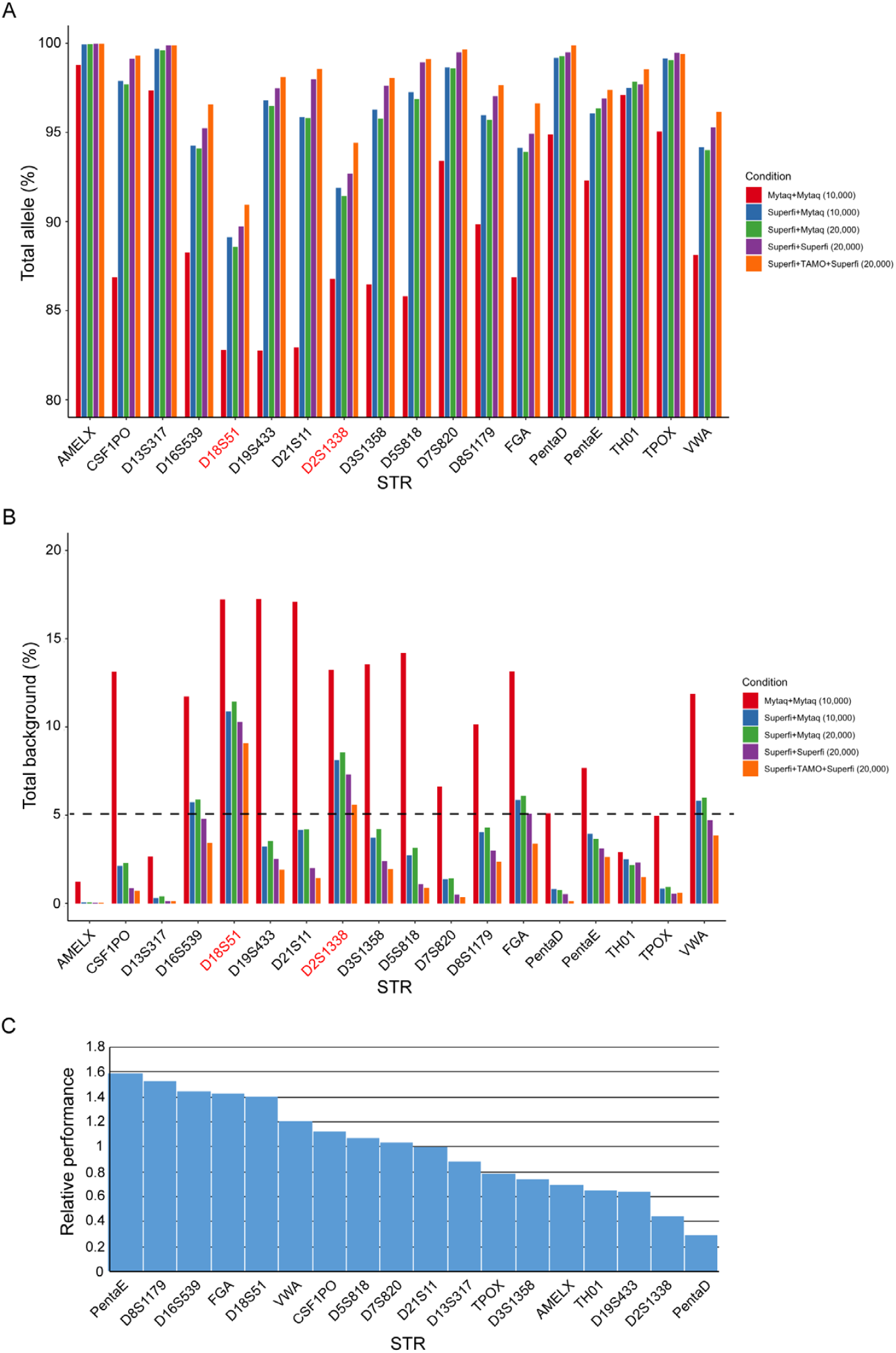
Human STR loci. (A) Percent of correct call on STR repeat length in each STR locus among different PCR conditions. Figure legend defines the polymerase used for each step of the NGS library construction (PCR1 and PCR2). The number of input cells is shown in parenthesis. (B) Percent of total background reads including stutter and noise in each STR locus among different PCR conditions. Underperforming STR loci with >5% background using the optimized PCR conditions are highlighted in red. (C) Relative STR locus performance (locus specific reads vs. mean of total reads).

**Figure S2.**
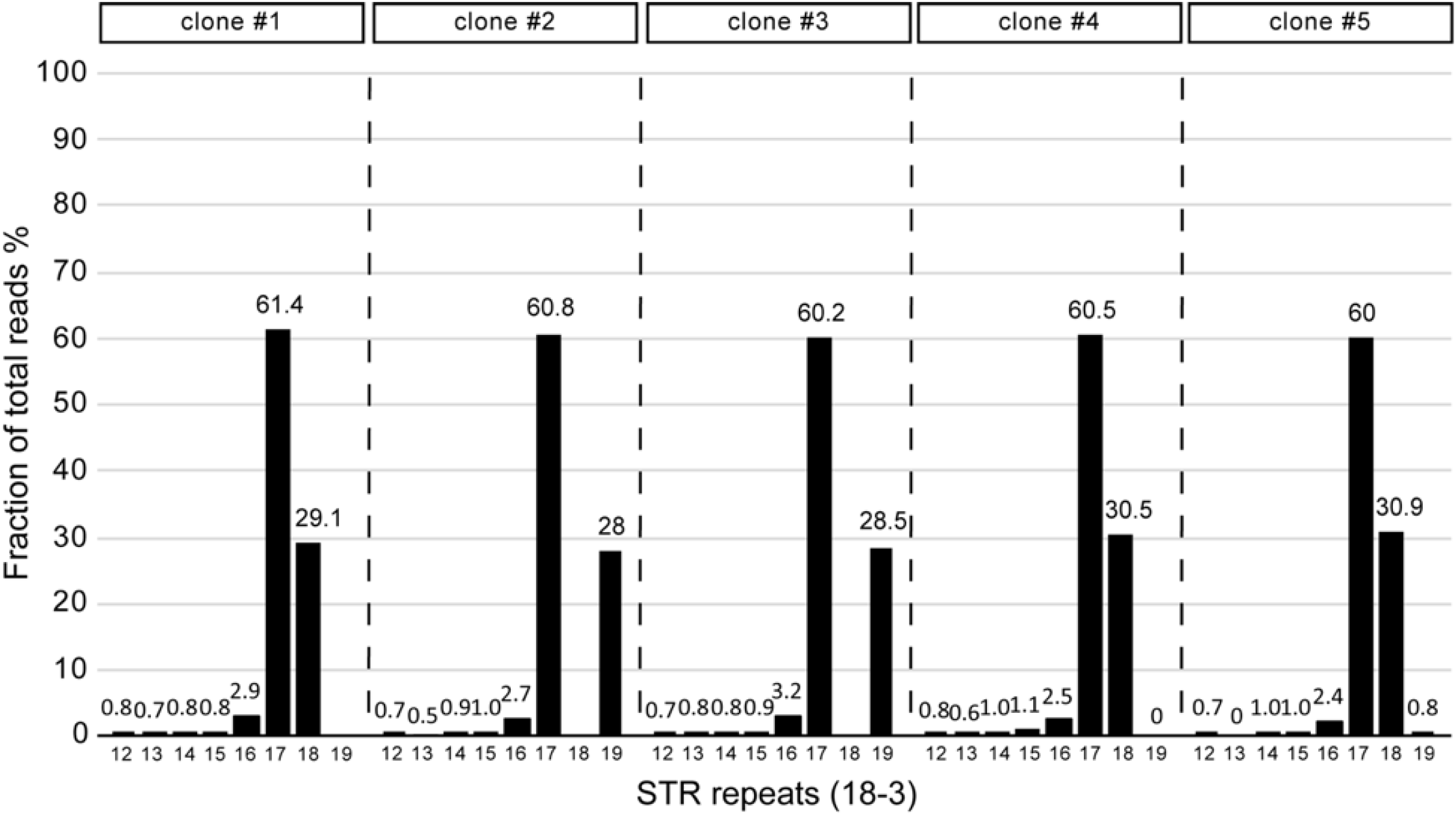
18-3 STR locus in NIH3T3 cells. Bar graph displays the percent of each STR repeat length of 18-3 locus in single cell clones of NIH3T3 cell line.

**Figure S3.**
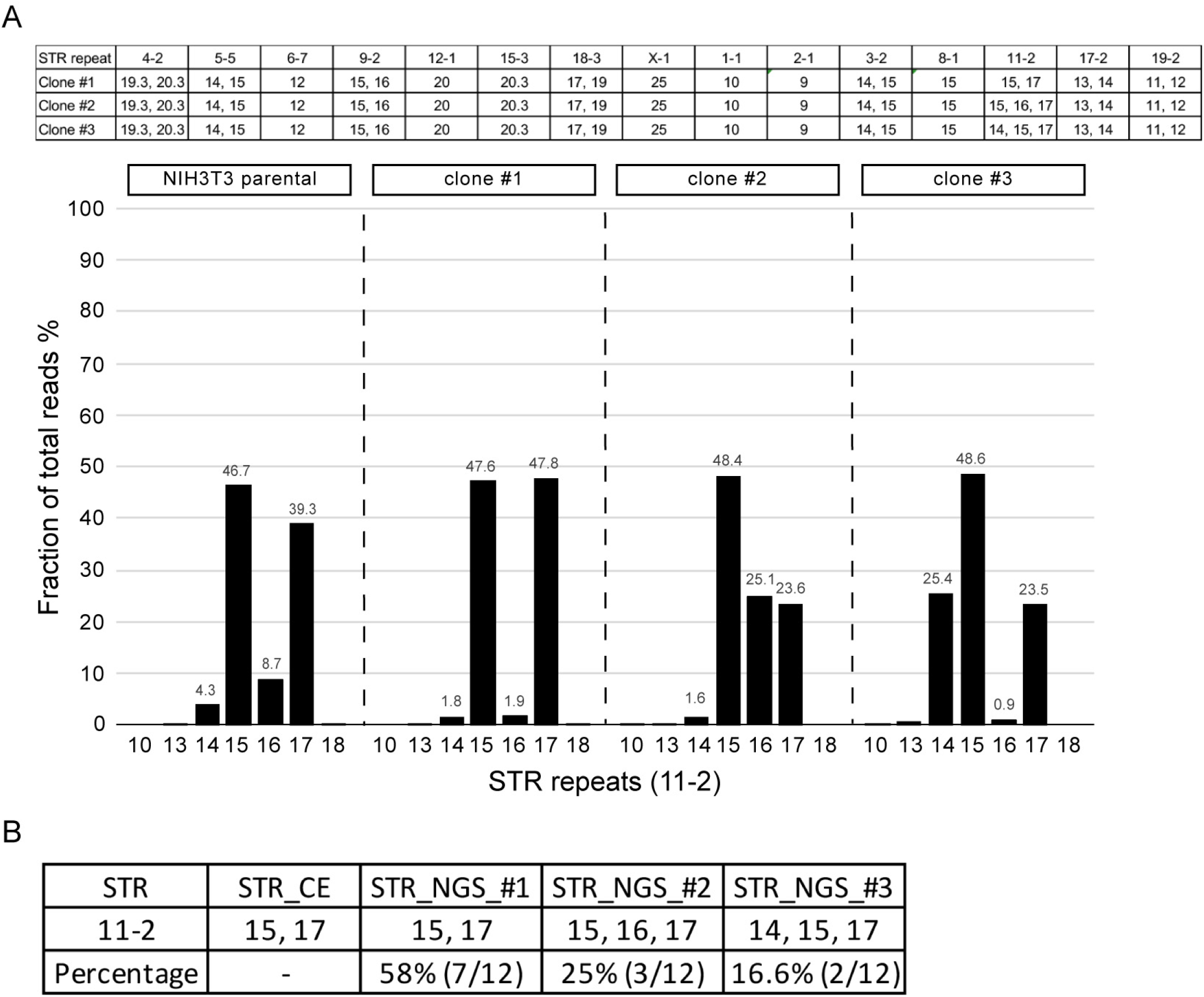
11-2 STR loci in NIH3T3 cells. (A) Table reports STR repeat lengths of 15 mouse STR loci in single cell clones representing three different subpopulations from NIH3T3 cells. Bar graph demonstrates the percent of each STR repeat length of 11-2 locus in three single cell clones and parental cells. (B) Proportion of three different populations reported for STR repeat at 11-2 locus.

**Figure S4.**
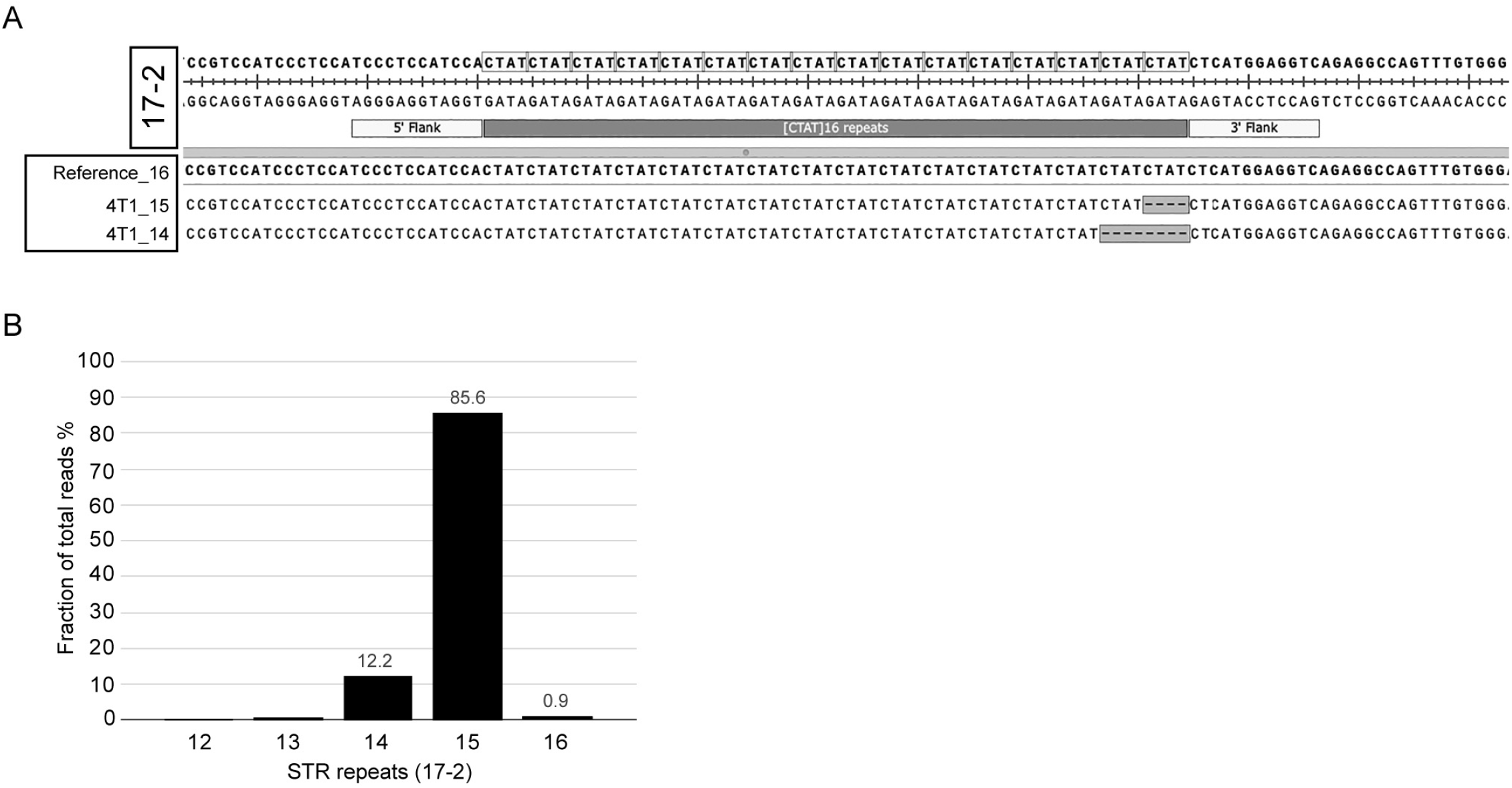
Sequencing alignment of 17-2 STR locus. (A) Sequence alignments of representative NGS reads among different STR repeat lengths for 17-2 STR locus of 4T1 cell line. (B) Percentage of each STR repeat length in 17-2 locus 4T1 cell line.

**Table S2.**
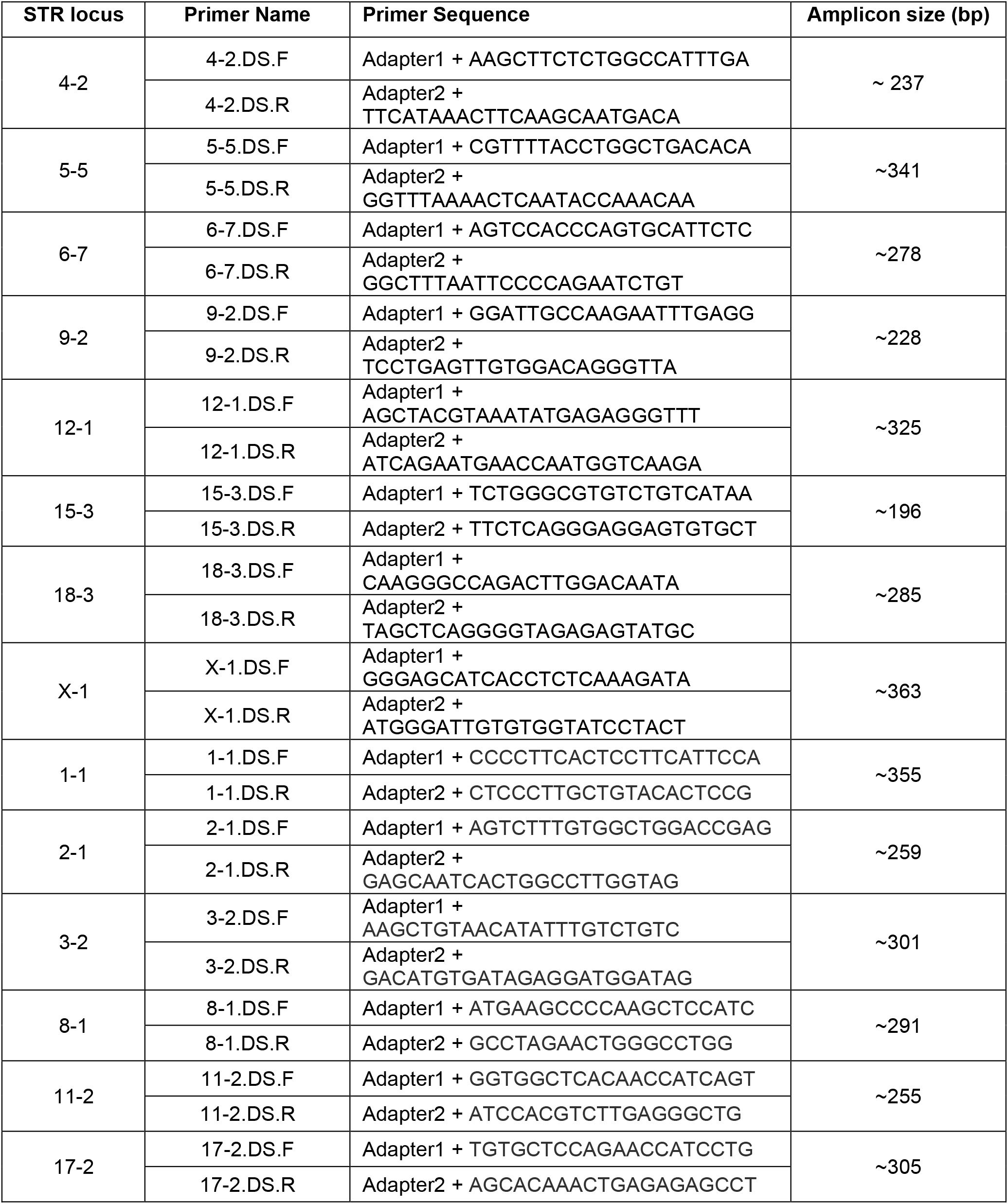
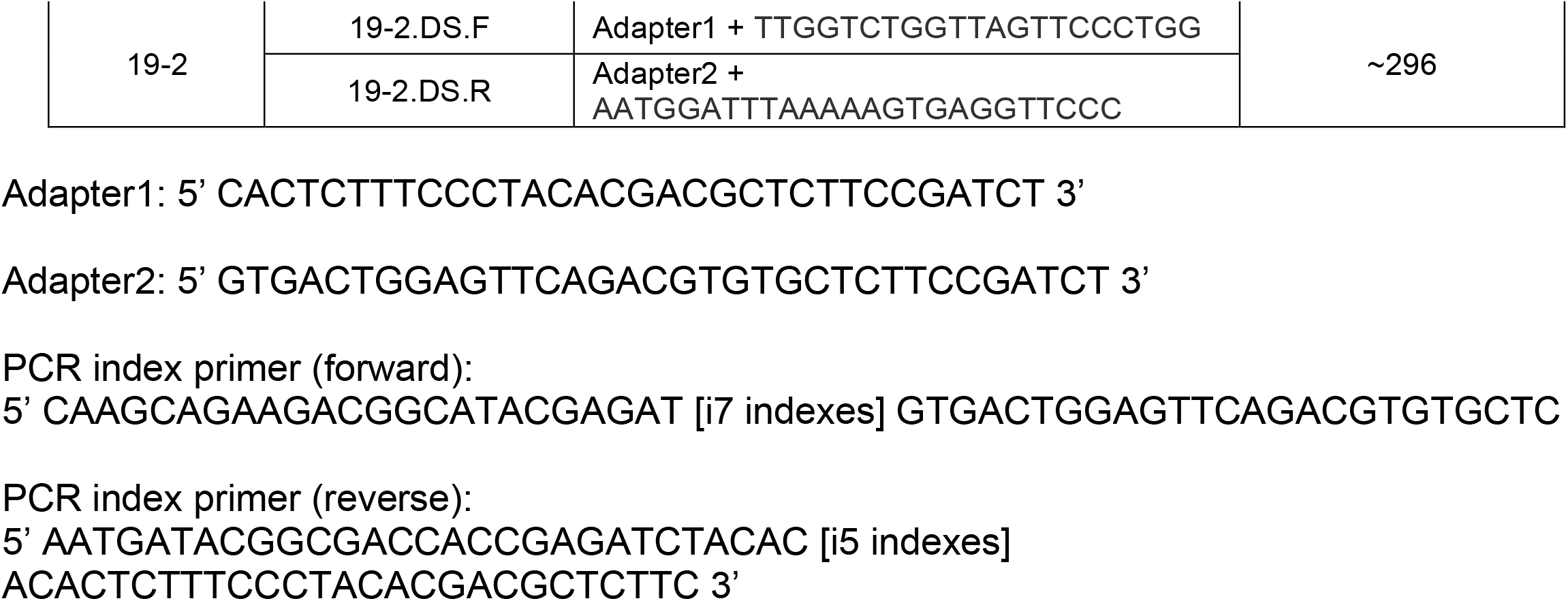
Mouse STR primer set.

**Table S1.**
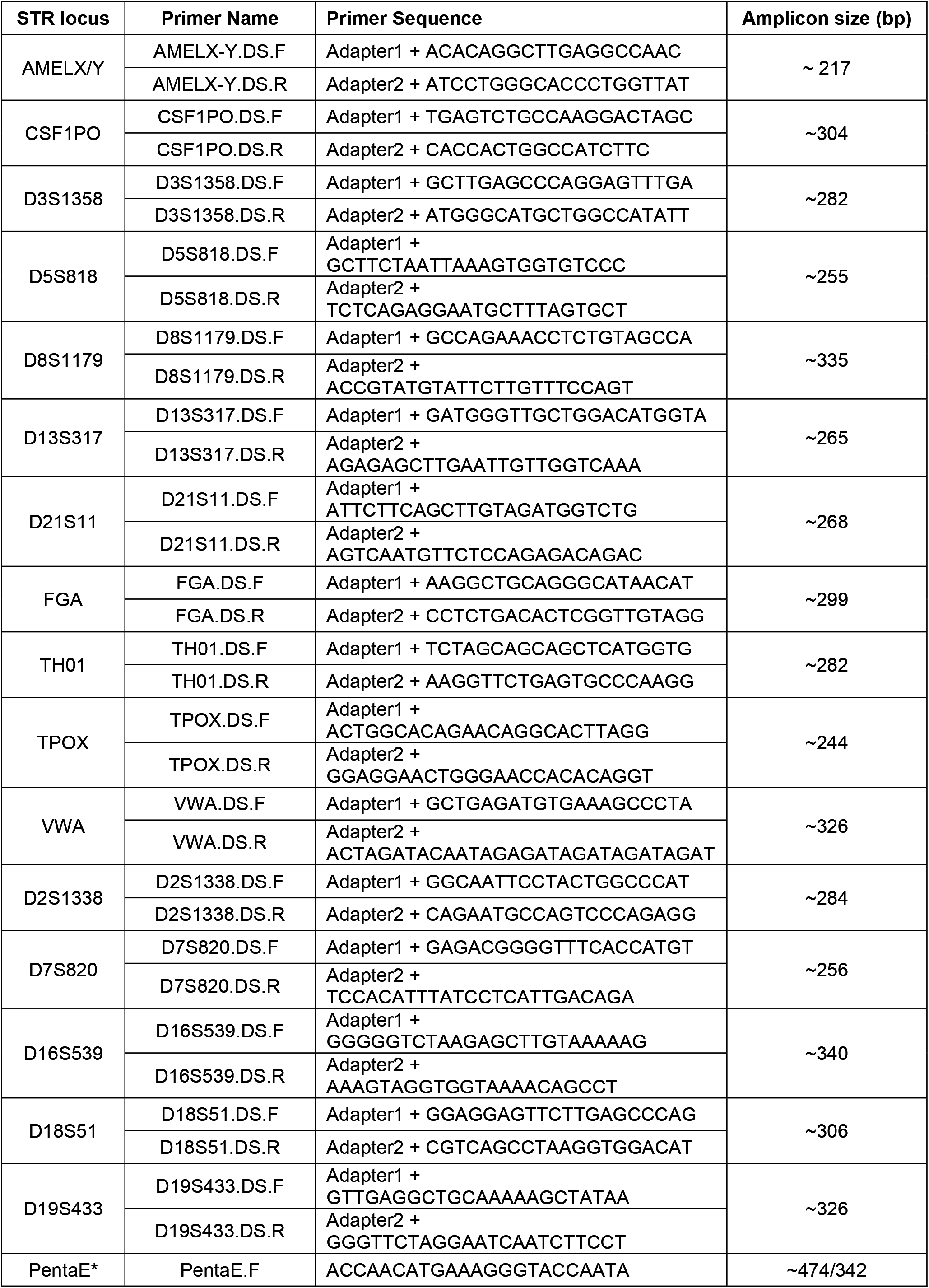
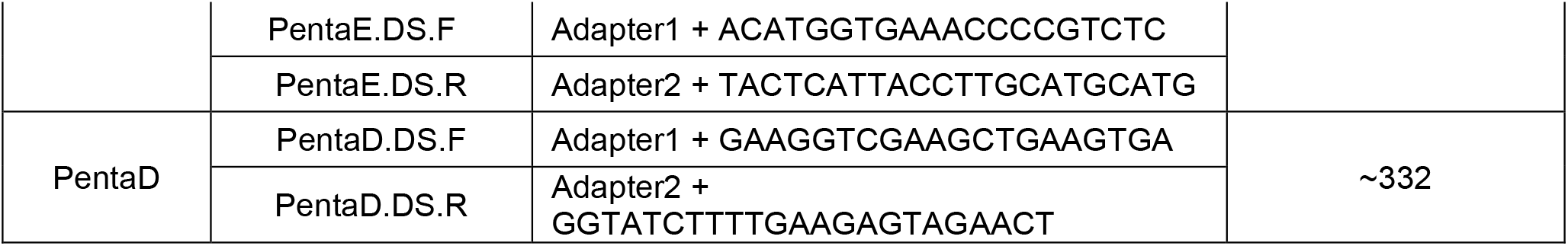
Human STR primer sets.

